# SLICER: Seamless Loss of Integrated Cassettes Using Endonuclease Cleavage and Recombination in *Deinococcus radiodurans*

**DOI:** 10.1101/2022.10.15.512367

**Authors:** Stephanie L. Brumwell, Katherine D. Van Belois, Daniel P. Nucifora, Bogumil J. Karas

## Abstract

Methods for creating seamless genome modifications are an essential part of the microbial genetic toolkit that allows for strain engineering through the recycling of selectable markers. Here, we report the development of a method, termed SLICER, which can be used to create seamless genome modifications in *D. radiodurans*. We used SLICER to sequentially target four putative restriction-modification (R-M) system genes, recycling the same selective and screening markers for each subsequent deletion. A fifth R-M gene was replaced by a selectable marker to create a final *D. radiodurans* strain with 5 of the 6 putative R-M systems deleted. While we observed no significant increase in transformation efficiency, SLICER is a promising method to obtain a fully restriction-minus strain and expand the synthetic biology applications of *D. radiodurans* including as an *in vivo* DNA assembly platform.

**GRAPHICAL ABSTRACT:** 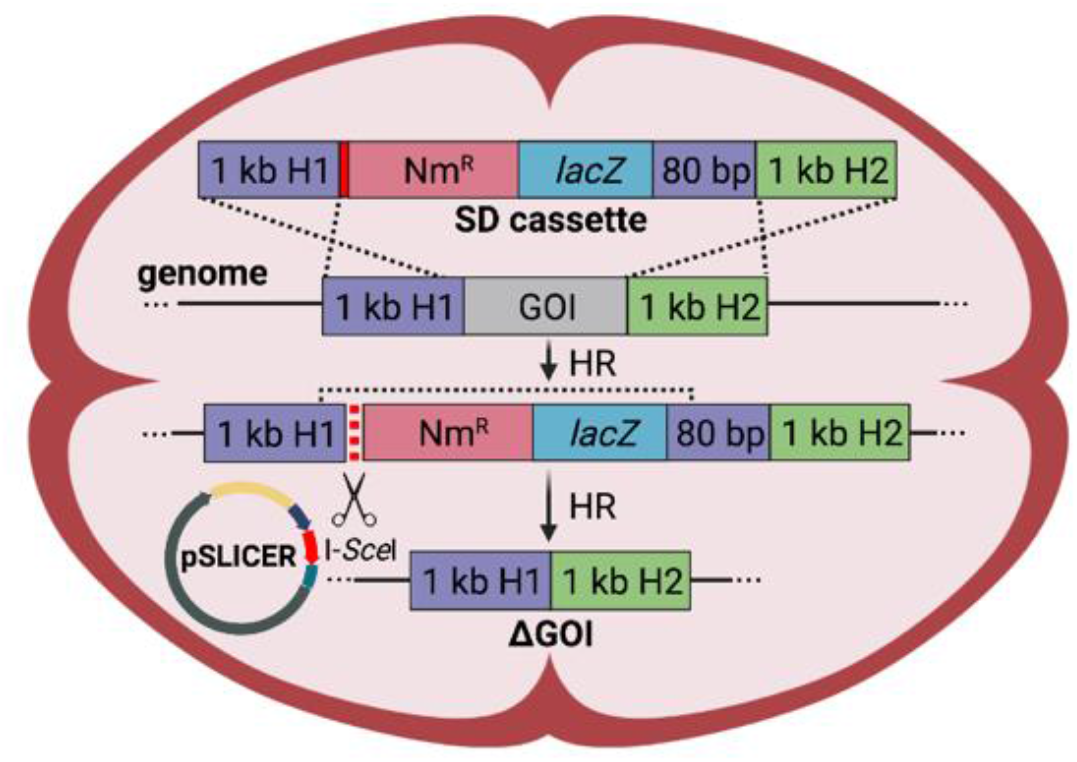

## INTRODUCTION

*Deinococcus radiodurans* is a polyextremophile bacterium with exceptional resistance to the lethal effects of ionizing and ultraviolet (UV) radiation, desiccation and other DNA-damaging agents^1,2^. This resistance has been linked to the superior homologous recombination and DNA repair mechanisms of this bacterium^3^, which have been shown to efficiently repair the genome as well as exogenous plasmid DNA following irradiation^4^. By exploiting this machinery, *D. radiodurans* has the potential to be a platform for microbial bioproduction, bioremediation, and synthetic biology applications^5–7^. This bacterium could act as a DNA assembly platform, complementing the most common method of assembling large DNA constructs and whole genomes in *Saccharomyces cerevisiae*^8,9^. Therefore, developing genetic tools for strain engineering and the study of *D. radiodurans* biology has become a priority.

Transformation and maintenance of synthetic constructs into the genome is most commonly achieved in *D. radiodurans* using antibiotics and their respective resistance gene. However, to make multiple seamless gene deletions in a single strain, it would be beneficial to have a method to recover selectable markers for reuse. Recently, a Cre-lox system was developed to allow for the removal of integrated selectable markers^10^. However, this method leaves behind *loxP* sites; therefore, there is currently no method for creating seamless gene deletions in this bacterium. To address this, we developed the SLICER method (seamless loss of integrated cassettes using endonuclease cleavage and recombination). One application for such a method would be for the generation of a restriction-minus strain.

Many microorganisms have restriction-modification (R-M) systems as part of the bacterial immune system, protecting against foreign DNA molecules^11^. Putative R-M systems in *D. radiodurans* R1 have been identified throughout the two chromosomes and two plasmids, which have been summarized on REBase (http://rebase.neb.com/, reference #22767)^12^. The genome contains four predicted Type II and two Type IV R-M systems containing restriction endonucleases, as well as a lone methyltransferase on the CP1 plasmid. Previous studies have characterized some of the R-M systems empirically^13–15^ and showed that they may be preventing the efficient transformation of *D. radiodurans*^16^.

Therefore, we used SLICER to sequentially delete four R-M systems in *D. radiodurans*. Replacement of the fifth R-M system with a neomycin selectable marker resulted in a strain lacking five of the six R-M systems. Deletion of all five systems did not affect bacterial growth and did not significantly improve transformation efficiency of the pRAD1 plasmid (6 kb). While transformation was not significantly improved in our final strain, the SLICER method was demonstrated as an efficient method for engineering *D. radiodurans* that will enable the deletion of multiple nonessential genes of interest (GOI) and ultimately lead to further development of laboratory or industrial strains.

## RESULTS AND DISCUSSION

We sought to develop a method for generating seamless deletions in the *D. radiodurans* genome. To achieve this, we modified the *S. cerevisiae* tandem repeat coupled with endonuclease cleavage (TREC) method^17^ to create the SLICER method for *D. radiodurans* engineering. A nonreplicating multi-host shuttle plasmid, termed the seamless deletion plasmid (pSD), was first built specifically for the targeted DNA region or GOI (Figure 1A). The key component of this plasmid, herein referred to as the seamless deletion (SD) cassette, contains a neomycin resistance gene and *lac*Z marker for selection and visual screening in *D. radiodurans*. These markers are flanked by two 1 kb regions homologous to the sequences upstream and downstream of the genomic target. Following homology region 1, there is an 18-bp I-*Sce*I endonuclease recognition site and prior to the second homology region there is a duplication of the last 80 bp of homology region 1.

**Figure 1.**
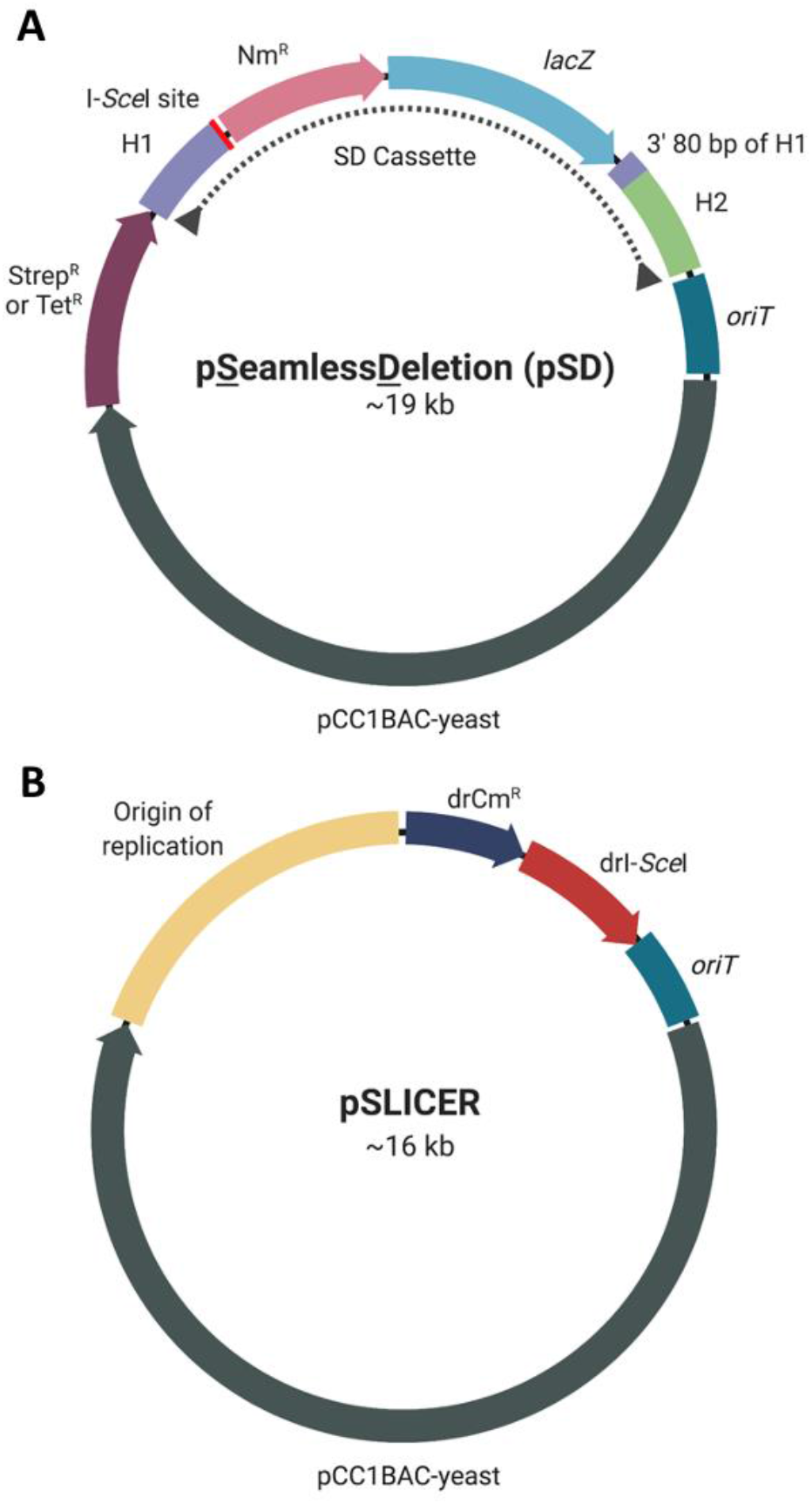
Plasmid maps of pSD and pSLICER. (A) Representative schematic of pSD illustrating the components contained on pSD1-pSD4: homology region 1 (H1), I-*Sce*I endonuclease recognition site, neomycin resistance gene (Nm^R^), β-galactosidase gene (*lacZ*), duplication of the 3’ 80 bp of H1, homology region 2 (H2), origin of transfer (*oriT*), pCC1BAC-yeast backbone for replication and selection in yeast and *E. coli*, streptomycin or tetracycline resistance gene (Strep^R^ or Tet^R^). The SD cassette is indicated with a dotted line. (B) Schematic of pSLICER containing an origin of replication for *D. radiodurans*, chloramphenicol resistance gene (drCm^R^), codon-optimized I-*Sce*I endonuclease (drI-*Sce*I). Created with BioRender.com.

An overview of the SLICER method is depicted in Figure 2. The first step is the integration of the SD cassette into the *D. radiodurans* genome at the target locus. The SD cassette was PCR amplified and delivered via chemical transformation into *D. radiodurans*. Alternatively, the whole pSD plasmid can be delivered via conjugation. Recombination between the two homology regions and their corresponding genomic regions results in integration of the SD cassette into the *D. radiodurans* genome, replacing the target GOI. *D. radiodurans* transformants containing the SD cassette are selected on TGY media supplemented with neomycin (5 μg mL^-1^) and X-Gal (40 μg mL^-1^) and appear blue in colour due to the expression of *lacZ*. The resulting strain is referred to as *D. radiodurans* + SD.

**Figure 2.**
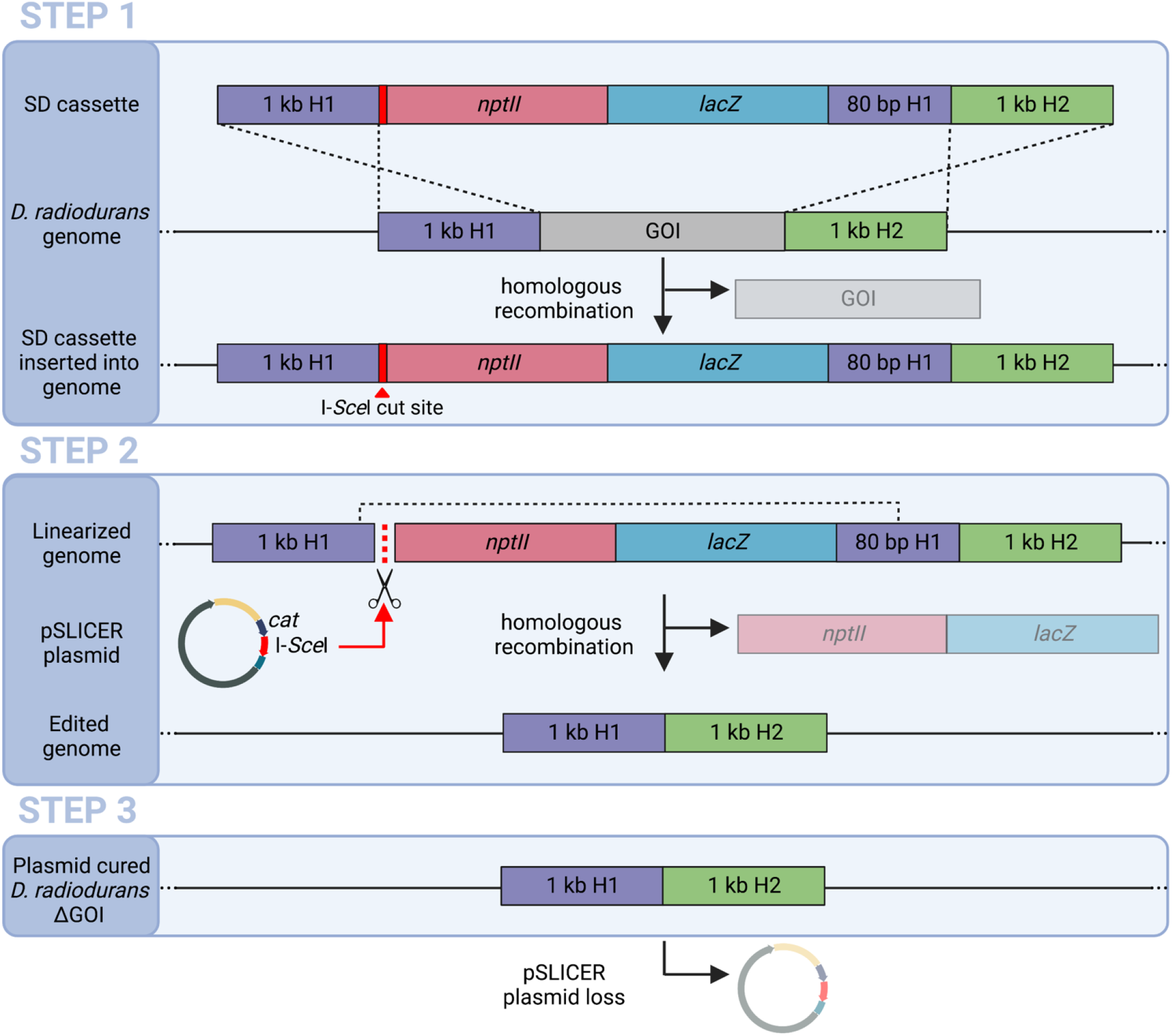
Overview of the SLICER method. STEP 1: Transformation of the SD cassette, containing *nptII* and *lacZ* genes for antibiotic selection and visual screening, into *D. radiodurans*. Homologous recombination of the 1 kb H1 and H2 regions occurs with the *D. radiodurans* genome resulting in integration of the SD cassette replacing the GOI. STEP 2: Conjugation of the pSLICER plasmid into *D. radiodurans* where it expresses the codon-optimized I-*Sce*I homing endonuclease that cuts at the 18-bp I-*Sce*I restriction site within the SD cassette. This double-strand break prompts a second homologous recombination event between the duplicated 3’ 80 bp of H1, removing the *nptII* and *lacZ* markers. STEP 3: Finally, plasmid curing to remove pSLICER results in a marker-free *D. radiodurans* ΔGOI strain. SD, seamless deletion; H1, homology region 1; H2, homology region 2; GOI, gene of interest; *nptII*, neomycin resistance gene; *lacZ*, β-galactosidase. Created with BioRender.com.

The second step in the SLICER method is the excision of the SD cassette facilitated by the pSLICER plasmid. We constructed the replicating helper plasmid, pSLICER, with a codon-optimized I-*Sce*I endonuclease, an origin of replication and a chloramphenicol selective marker for *D. radiodurans* (Figure 1). The I-*Sce*I enzyme was chosen because there are no recognition sites present in the wild-type genome of *D. radiodurans*. The I-*Sce*I endonuclease was designed under the regulation of the PDR_2508 promoter and terminator set as it was shown to have high expression in *D. radiodurans* but low expression in *E. coli*^18^. This plasmid was then transformed into an *E. coli* Δ*dap*A strain harbouring the conjugative plasmid pTA-Mob^19^. Conjugation of pSLICER from the *E. coli* conjugative donor strain to *D. radiodurans* + SD was then performed. The I-*Sce*I endonuclease produces a double-stranded break at the I-*Sce*I recognition sequence within the SD cassette, stimulating homologous recombination between homology region 1 and the 80-bp duplicated region. This recombination event excises the two markers. Transconjugants are selected on TGY media supplemented with chloramphenicol (3 μg mL^-1^) and X-Gal. Contrary to the screening in Step 1, transconjugants that have had the SD cassette excised should appear pink since they have lost the *lac*Z gene. The resulting strain is referred to as *D. radiodurans* + SLICER. The codon-optimized I-*Sce*I endonuclease was proven to be functional in *D. radiodurans* and essential for SD cassette excision and ultimately for the success of the SLICER method (Supplemental Note, Figure S1).

The final step in the SLICER method is to cure pSLICER from the seamless deletion strain. The *D. radiodurans* + SLICER strain was grown in nonselective media overnight and dilutions were subsequently spot plated on nonselective media. Resulting single colonies were then struck on nonselective media as well as media supplemented with either chloramphenicol or neomycin. The colonies are confirmed to be cured of the plasmid when growth is observed on nonselective plates but not on selective plates. At the end of the seamless deletion process, the resulting *D. radiodurans* ΔGOI strain will have the target gene deleted with no remnants of the process remaining in the genome or the cell. The entire SLICER method can be completed in approximately two weeks and the step-by-step protocol is summarized in Figure S2.

Using the seamless deletion strategy outlined above, we performed the sequential deletion of four R-M system genes in the *D. radiodurans* genome (Figure 3), with the fifth R-M system (ORF2230) subsequently deleted using homologous recombination-based integration of a neomycin marker. Four nonreplicating pSD plasmids, named pSD1-pSD4, were built for each target R-M gene: ORF14075, *Mrr*, ORF15360, and *Mrr2*, which will herein be called RM1, RM2, RM3 and RM4, respectively. These target genes were named numerically in the order that they were used to generate deletions. Each plasmid contains the same elements apart from the homology regions, which are specific to each target gene (Figure 1).

**Figure 3.**
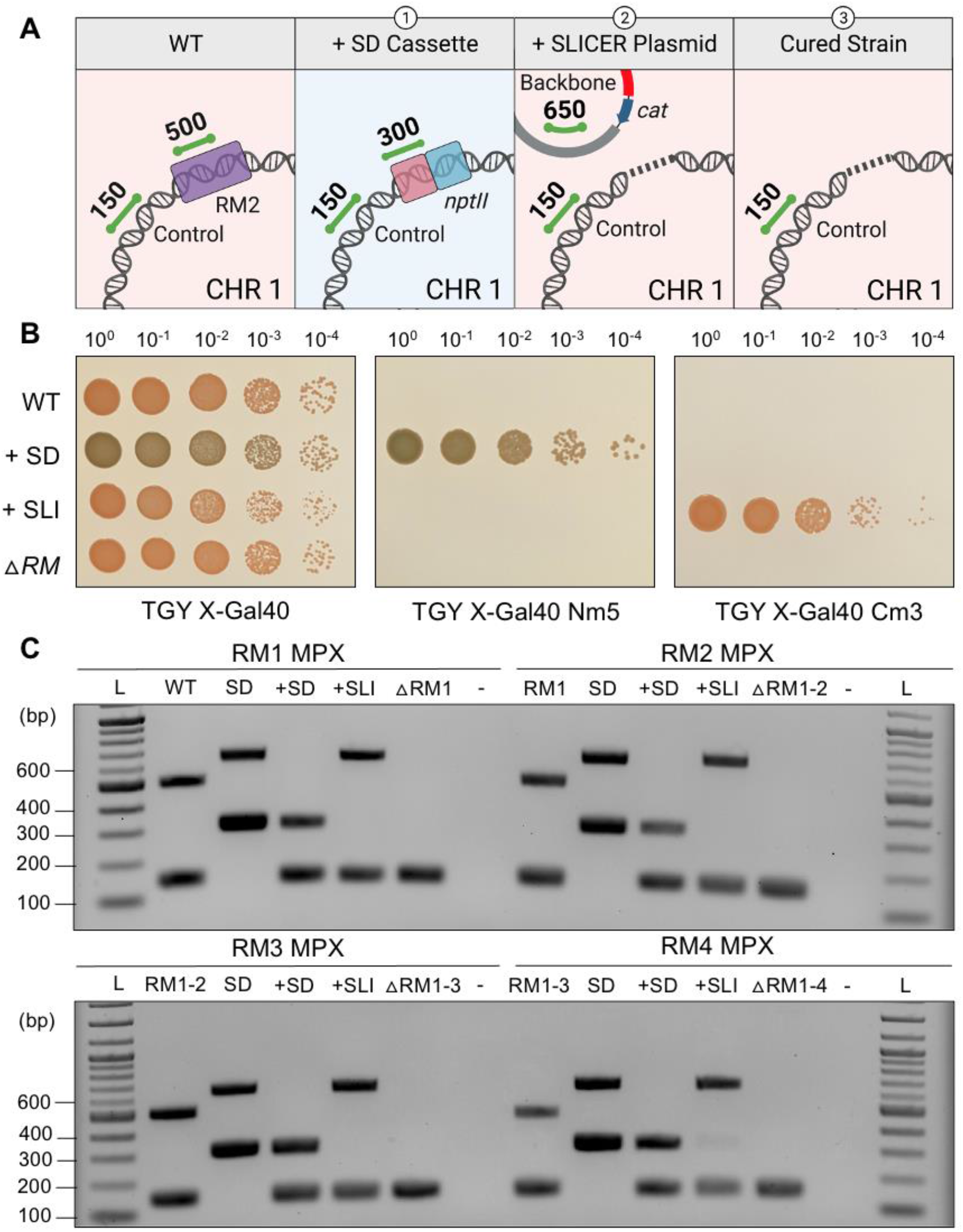
Seamless deletion of RM1-4 genes using SLICER in *D. radiodurans*. (A) Representative schematic of the multiplex PCR amplicons present in *D. radiodurans* strains: 1) wild type (WT), 2) following integration of the SD cassette at the RM locus (+SD), 3) following conjugation of pSLICER and excision of the SD cassette (+SLI), and 4) following curing of pSLICER curing (ΔRM). Expected multiplex PCR amplicons are shown as green lines with the corresponding size in bp. Created with BioRender.com. (B) Spot plates of 10-fold serial dilutions of the same strains listed in (A). All plates contain X-Gal 40 μg mL^-1^. (C) Gel electrophoresis of multiplex PCR analysis (RM1-RM4 MPX) of a single *D. radiodurans* colony from each step in the creation of the four seamless R-M gene deletions in the order depicted in (A): WT, +SD +SLI, and following plasmid curing (ΔRM1, ΔRM1-2, ΔRM1-3, and ΔRM1-4). Additional controls include the SD plasmid DNA extracted from *E. coli* (SD). Expected amplicon sizes are approximately 150 bp for the *D. radiodurans* gDNA control, 300 bp for *nptII* in the SD cassette, 500 bp for the R-M gene, and 650 bp for the pSLICER backbone. L, 1 kb plus ladder.

Following Step 1, 2 and 3 of the SLICER method (Figure 2), the *D. radiodurans* genome was analyzed to confirm insertion of the SD cassette, removal of the SD cassette, and curing of the pSLICER plasmid. Gene deletion analysis of RM1, RM2, RM3 and RM4 is shown in Figure 3 resulting in the creation of *D. radiodurans* ΔRM1, ΔRM1-2, ΔRM1-3 and ΔRM1-4 strains, respectively. Analysis was conducted by spot plating dilutions on nonselective media and media supplemented with neomycin or chloramphenicol, all of which contained X-Gal (Figure 3B). In addition, multiplex PCR analysis was performed on DNA extracted from one individual colony for each seamless deletion event (RM1-4) (Figure 3C). If present in the examined DNA, the multiplex PCR should amplify a 150 bp amplicon at a non-target site in the *D. radiodurans* genome, a 300 bp amplicon within the neomycin marker on the SD cassette, a 500 bp amplicon within the target gene (RM1-4), and/or a 650 bp amplicon within the pSLICER backbone. The position and size of the expected amplicons following each step of the seamless deletion strategy are depicted in Figure 3A.

From the analyses of the RM1-RM4 deletions (Figure 3B,C), we observed that the wild-type *D. radiodurans* strain was only able to grow on nonselective media, appeared pink in colour, and the PCR results showed amplification of the genomic DNA control and target gene. Multiplex PCR performed on the pSD1-4 plasmids containing the SD cassette showed amplification of the neomycin marker and backbone amplicons. Following integration of the SD cassette, *D. radiodurans* + SD was able to grow on the nonselective and neomycin-supplemented media, and appeared blue in colour on both. The PCR results showed amplification of the genomic control and notably, there was no amplification of the target gene amplicon. Rapid gene deletions were observed across all genomic copies of *D. radiodurans* using this method (after a single passage on selective media) compared to previous methods that reported the need to subculture for 30-35 passages alternating growth in liquid and solid media to obtain homozygous gene knockouts ^20^.

After conjugating in the pSLICER plasmid, *D. radiodurans* + SLICER is able to grow on nonselective and chloramphenicol-supplemented media, but not neomycin-supplemented media. With the loss of the SD cassette, the colonies once again appeared pink. The PCR results showed amplification of the genomic control and backbone amplicons. Notably, there is no amplification of the neomycin marker. Finally, curing of the pSLICER plasmid from the *D. radiodurans* ΔRM strain only allows for growth on the nonselective plate as the strain no longer contains the pSLICER plasmid, and colonies still appear pink. Multiplex PCR only shows amplification of the genomic control amplicon, indicating that the pSLICER plasmid was successfully cured.

Following the fourth deletion, the fifth R-M system was deleted using homologous-recombination based integration of a neomycin marker using the cassette from pDEINO10 previously used to delete ORF2230^21^. The final *D. radiodurans* ΔRM1-5 Nm^R^ strain can be propagated with neomycin selection (Figure 4A). Further confirmation that all four genes (RM1-4) have been seamlessly deleted in the *D. radiodurans* ΔRM1-4 strain was performed using multiplex PCR (Figure 4B). The sixth R-M system that has not yet been deleted is *mcrBC* on the MP1 megaplasmid. A *D. radiodurans* mutant was previously created with an insertion in the *mcrB* gene, which did not lead to an increase in transformation efficiency^16^. The low transformation efficiency in *D. radiodurans* may be the result of multiple active R-M systems; therefore, it is unlikely that an improvement would be observed by deleting a single system.

**Figure 4.**
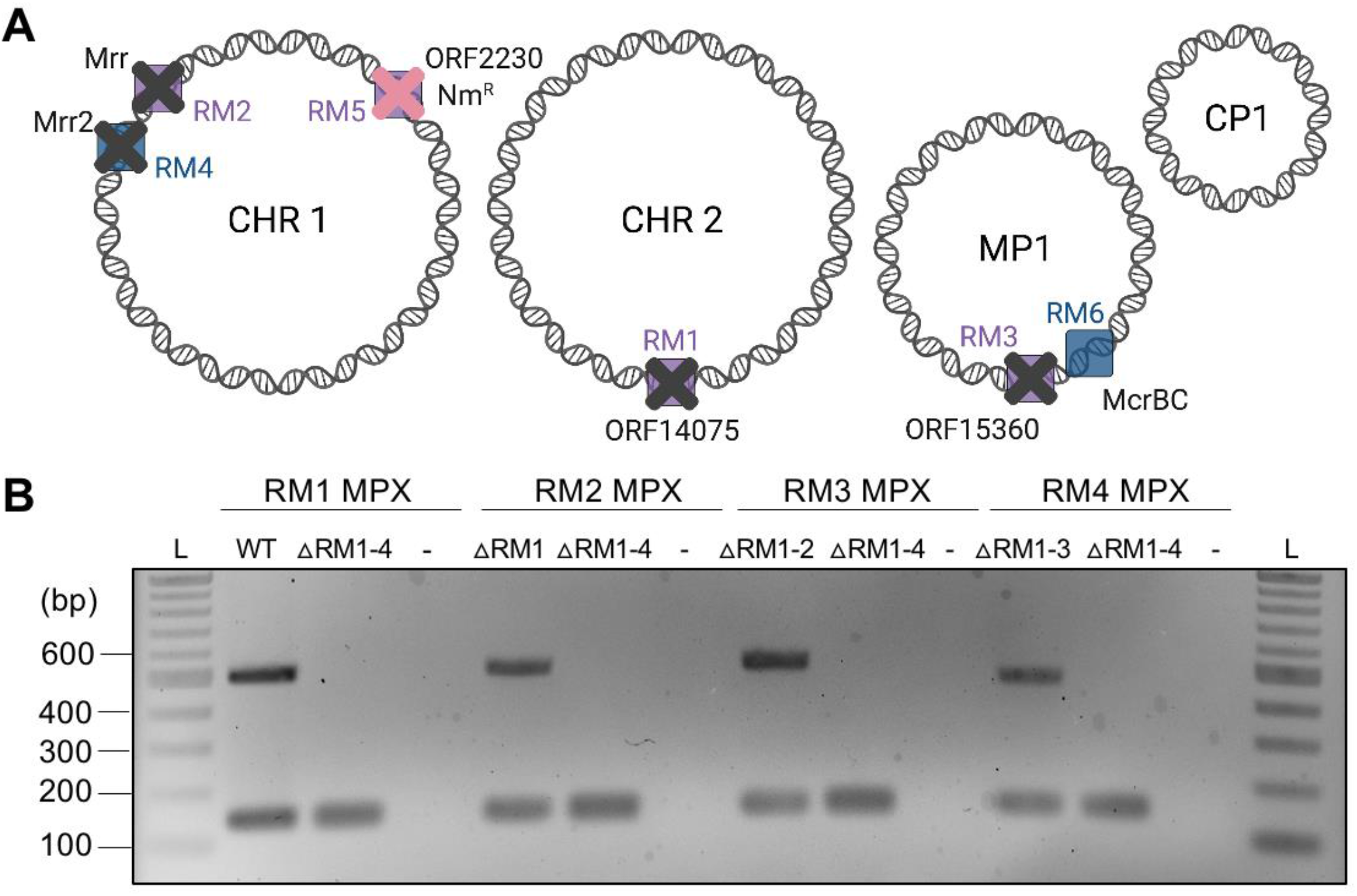
*D. radiodurans* ΔRM1-4 multiplex PCR analysis. (A) Schematic representation of the *D. radiodurans* ΔRM1-5 Nm^R^ genome with the first four R-M genes (RM1, RM2, RM3, RM4) seamlessly deleted as indicated by grey X’s, and the fifth R-M system (RM5) replaced with a neomycin marker (Nm^R^) as indicated by a pink X. Created with BioRender.com. (B) Gel electrophoresis of four multiplex PCR analyses (RM1–RM4 MPX) for each seamless gene deletion performed on a single *D. radiodurans* ΔRM1-4 colony. For the RM1 multiplex, a *D. radiodurans* wild-type (WT) genomic DNA control is used, and for all subsequent multiplex analyses the cured strain from the previous deletion was used as a control (ΔRM1, ΔRM1-2, and ΔRM1-3, respectively). A negative control (-) where water was used in place of template was also included. Expected amplicon sizes are approximately 150 bp for the *D. radiodurans* gDNA control and 500 bp for the R-M gene, if present. L, 1 kb plus ladder.

Physiological analysis of *D. radiodurans* ΔRM strains was performed by testing their growth in liquid TGY media. The growth phenotype of *D. radiodurans* ΔRM strains compared to wild type revealed no significant difference based on the growth curve, endpoint density or calculated growth rates (Figure 5A). This suggests that removal of the first five R-M system genes did not result in any growth deficits in *D. radiodurans*, which is promising if this strain (or the full restriction minus strain) is to be used as synthetic biology chassis or for microbial bioproduction.

**Figure 5.**
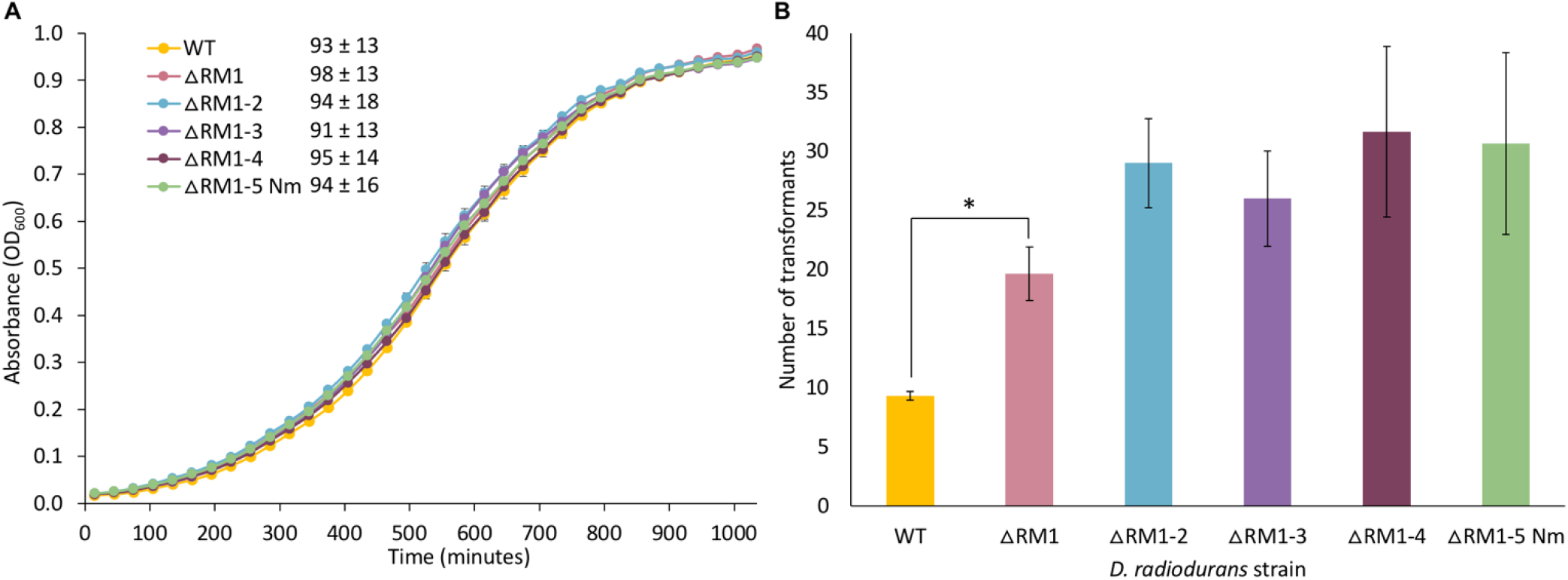
Physiological analysis of *D. radiodurans* R-M deletion strains. (A) Growth curves of *D. radiodurans* WT and ΔRM knockout strains grown in liquid TGY media for 17 hours. Each data point represents the mean of three biological and two technical replicates, with error bars representing standard error of the mean. The doubling time for each strain is reported (in minutes) in the legend and represents the mean value of the three biological and two technical replicates ± the standard deviation. (B) Number of *D. radiodurans* transformants that grew on TGY media supplemented with chloramphenicol 3 μg mL^-1^ following heat shock transformation of the pRAD1 plasmid to *D. radiodurans* WT and ΔRM knockout strains. 850 ng of DNA was used for each transformation. The data is presented as a bar graph displaying the mean of three biological replicates with error bars representing standard error of the mean. A two-sample, one-tail student’s t-test was used to compare WT (control) to each ΔRM knockout strain: *P<0.05.

Transformation of *D. radiodurans* ΔRM strains was performed using the ∼6 kb pRAD1 plasmid to determine if these strains have higher transformation efficiency compared to wild type. Heat shock transformation was performed using 850 ng of plasmid DNA isolated from *E. coli* Epi300 into wild type and all five ΔRM strains (Figure 5B). We observed an average transformation efficiency of 3.86 × 10^1^ and 1.27 × 10^2^ CFU/μg DNA for wild type and *D. radiodurans* ΔRM1-5 Nm^R^, respectively. These results indicate that through the deletion of five R-M systems, we are able to achieve a higher number of transformants on average but there is no significant difference in transformation efficiency compared to wild type.

In summary, we have created the first seamless genome modification strategy for *D. radiodurans* engineering and demonstrated that the SLICER method can be used for the sequential deletion of endogenous genes. Using this method, homozygous deletions can be made rapidly across all copies of the *D. radiodurans* genome, and it is the first report of the *I-Sce*I endonuclease being used in this bacterium. The SLICER method developed here should enable seamless deletion of the remaining R-M system(s), and any nonessential DNA region of interest in *D. radiodurans* such as those involved in the biosynthesis of essential amino acids for the generation of auxotrophic strains, or proteases in bioproduction strains. The deletion of five of the six known restriction-systems in *D. radiodurans* is a major step towards the creation of a fully restriction minus strain, which we hypothesize will significantly improve transformation of DNA to *D. radiodurans*. The development of a restriction minus strain will expand the synthetic biology applications of *D. radiodurans* as a host for DNA assembly and may allow for genome reduction or replacement for the study of extremophile biology.

## METHODS

### Microbial Strains and Growth Conditions

*Deinococcus radiodurans* R1 was grown at 30°C in TGY medium (5 g L^− 1^ tryptone, 3 g L^−1^ yeast extract, 1 g L^−1^ potassium phosphate dibasic, and 2.5 mL of 40% w/v glucose) supplemented with appropriate antibiotics (chloramphenicol, 5 μg mL^−1^; neomycin, 5 μg mL^−1^). *Escherichia coli* (Epi300, Lucigen) was grown at 37°C in Luria Broth (LB) supplemented with chloramphenicol, 15 μg mL^−1^. *Escherichia coli* ECGE101 (Δ*dapA*)^22^ was grown at 37°C in LB media supplemented with DAP, 60 μg mL^−1^, and appropriate antibiotics (chloramphenicol, 15 μg mL^−1^; gentamicin, 40 μg mL^−1^). *Saccharomyces cerevisiae* VL6-48 (ATCC MYA-3666: MATα *his3*-Δ200 *trp1*-Δ1 *ura3*–52 *lys2ade2*–1 *met14 cir*^*0*^) was grown at 30°C in 2X YPAD rich medium (20 g L^−1^ yeast extract, 40 g L^−1^ peptone, 40 g L^−1^ glucose, and 80 mg L^−1^ adenine hemisulfate), or in complete minimal medium lacking histidine supplemented with 60 mg L^−1^ adenine sulfate (Teknova Inc.) with 1 M sorbitol. All strains created in this study are summarized in Supplemental Table D-1.

*Deinococcus radiodurans* R1 was grown at 30°C in TGY medium (5 g L^-1^ tryptone, 3 g L^-1^ yeast extract, 1 g L^-1^ potassium phosphate dibasic, and 2.5 mL 40% w/v glucose) supplemented with appropriate antibiotics (chloramphenicol 5 μg mL^-1^, neomycin 5 μg mL^-1^, tetracycline 0.5 μg mL^-1^). *Escherichia coli* (Epi300, Lucigen) was grown at 37°C in Luria Broth (LB) supplemented with chloramphenicol 15 μg mL^-1^. *Escherichia coli* ECGE101 (Δ*dapA*)^22^ was grown at 37°C in LB media supplemented with diaminopimelic acid (DAP) 60 μg mL^-1^ and appropriate antibiotics (chloramphenicol 15 μg mL^-1^ and/or gentamicin 40 μg mL^-1^). *Saccharomyces cerevisiae* VL6−48 (ATCC MYA-3666: MATα his3-Δ200 trp1-Δ1 ura3−52 lys2 ade2−1 met14 cir^0^) was grown at 30°C in 2X YPAD rich medium (20 g L^-1^ yeast extract, 40 g L^-1^ peptone, 40 g L^-1^ glucose, and 80 mg L^-1^ adenine hemisulfate), or in complete minimal medium lacking histidine supplemented with 60 mg L^-1^ adenine sulfate (Teknova Inc.) with 1 M sorbitol.

### CaCl2 Transformation of *D. radiodurans*

*For competent cells:* A 50 mL culture of *D. radiodurans* was grown to an OD600 of 0.2. The culture was transferred to a 50 mL falcon tube and centrifuged at 3000 *g* for 15 min at 4°C. The supernatant was discarded and the pellet was resuspended in 250 μL of ice-cold 0.1 M CaCl2 15% glycerol solution using gentle agitation. The competent cells were aliquoted in 50 μL increments, frozen in a −80°C ethanol bath and stored at - 80°C.

#### For transformation

50 μL of competent cells per reaction were thawed on ice for 15 min. Then, 5 μL of transforming DNA (linear PCR product or plasmid) was mixed with the competent cells. The mixture was incubated on ice for 30 min then heat shocked in a 42°C water bath for 45 seconds. The tubes were returned to ice for 1 min and 1 mL of 2X TGY media was added to each tube. The recovery cultures were transferred to a 50 mL falcon tube and grown with shaking at 30°C for 2 hours at 225 rpm. Finally, 300 μL of the transformation mixture was plated on TGY media with appropriate supplements (chloramphenicol 3 μg mL^-1^ or neomycin 5 μg mL^-1^ and/or X-Gal 40 μg mL^-1^) and incubated at 30°C for 2-3 days. Colonies were counted manually.

### Conjugation from *E. coli* to *D. radiodurans*

Conjugation from *E. coli* to *D. radiodurans* was performed as previously described ^21^, with the following modifications. The donor strain was *E. coli* ECGE101 Δ*dapA*^22^ harbouring pTA-Mob ^19^ and pSLICER. The *D. radiodurans* R1 recipient strains with the integrated SD cassettes were grown in TGY media supplemented with neomycin (5 μg mL^−1^) prior to conjugation. The selective plates were TGY media supplemented with chloramphenicol (3 μg mL^−1^).

## ASSOCIATED CONTENT

### Supporting Information

The Supplementary File contains Supplementary Note, Figures S1-S2, Tables S1-S3 and Supplementary Methods.

## AUTHOR INFORMATION

### Author Contributions

S.L.B., and B.J.K conceived the experiments, S.L.B, K.D.V.B, and D.P.N conducted the experiments, S.L.B, and B.J.K analyzed the results, S.L.B. wrote the paper, S.L.B. and B.J.K edited the final version of the paper. All authors have given approval to the final version of the manuscript.

### Notes

The authors declare no competing financial interests.

## ACKNOWLEDGMENT

This research was funded by Natural Sciences and Engineering Research Council of Canada (NSERC), grant number: RGPIN-2018-06172 (to B.J.K.). In addition to the figures noted, the graphical abstract was also created with BioRender.com.

## Notes

### Competing Interest Statement

The authors have declared no competing interest.

