## Supplemental file for "SLICER: Seamless Loss of Integrated Cassettes Using Endonuclease Cleavage and Recombination in *Deinococcus radiodurans*"

### Supplemental Note

We sought to verify that the codon-optimized I-*Sce*I endonuclease encoded on the pSLICER plasmid was not only functional in *D. radiodurans* but is necessary for the success of the SLICER method. To determine the frequency of SD cassette loss from the *D. radiodurans* genome following integration, dilutions of *D. radiodurans* ΔRM1-4 Nm$^R$, harbouring the SD cassette, were plated on nonselective and selective media (Figure S1A). Due to the presence of *lacZ* in the SD cassette, blue colonies should be indicative of those carrying the SD cassette while pink colonies indicate loss of the cassette. The percentage of *D. radiodurans* colonies that appeared pink with and without antibiotic selection were 1.1% and 2.1%, respectively, indicating the occurrence of natural SD cassette loss or mutation following propagation. The pink colonies obtained from both nonselective and selective plates were further analyzed by streaking them onto selective media (data not shown). All colonies were able to grow on selective media, indicating that while these colonies appeared to have lost or mutated the *lacZ* gene, the neomycin marker in the SD cassette was still functional. As such, the integrated SD cassettes appear to be quite stable and spontaneous loss of these cassettes could not be easily obtained by growing cultures without selective pressure.

As further confirmation that the I-*Sce*I endonuclease is required for excision of the SD cassette in the SLICER method, conjugation of pSLICER and a control plasmid, pDEINO1, was performed to *D. radiodurans* ΔRM1-4 Nm$^R$ with the integrated SD cassette (Figure S1B). The pDEINO1 plasmid contains all of the same components as pSLICER including a *D. radiodurans* origin of replication and chloramphenicol marker, but lacks the I-*Sce*I endonuclease. When this plasmid was conjugated to *D. radiodurans*, all transconjugant colonies appeared blue, indicating that they still harbored the SD cassette. Conversely, when the pSLICER plasmid was conjugated to *D. radiodurans*, all transconjugant colonies appeared pink, indicating that the SD cassette had been lost. This allowed us to conclude that the codon-optimized I-*Sce*I endonuclease is functional in *D. radiodurans* and is essential for SD cassette excision.

### Supplemental Figures

**A**

| Dilution | Nonselective | | Selective (Nm5) | |
|---|---|---|---|---|
| | Blue | Pink | Blue | Pink |
| $10^{-2}$ | TNTC | TNTC | 366 | 4 |
| $10^{-3}$ | 959 | 20 | 29 | 0 |
| $10^{-4}$ | 104 | 0 | 4 | 0 |
| **Percent** | | **2.04%** | | **1.08%** |

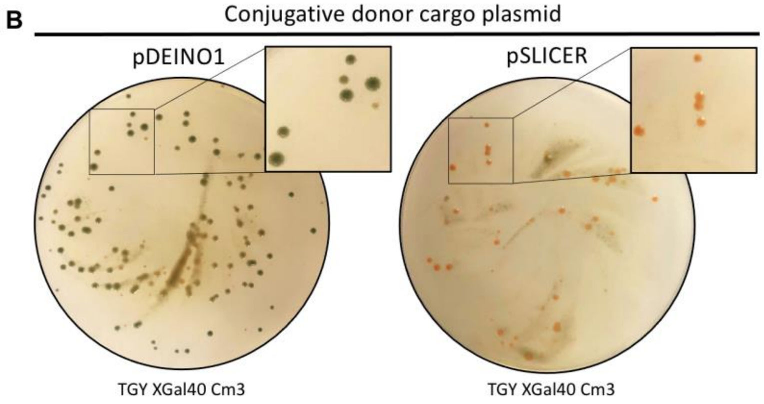


**Figure S1.** Validation of I-*Sce*I endonuclease function. (A) Serial dilutions of *D. radiodurans* ΔRM1-4 Nm$^R$ plated on nonselective (TGY X-Gal) and selective (TGY X-Gal supplemented with neomycin) media. The number of blue and pink colonies was counted and the percentage of colonies that appeared pink over total colonies is reported. (B) Selective plates following conjugation of pDEINO1 and pSLICER from *E. coli* to *D. radiodurans* $\Delta$RM1-4 Nm$^{R}$. Antibiotic concentrations are reported as μg mL$^{-1}$.

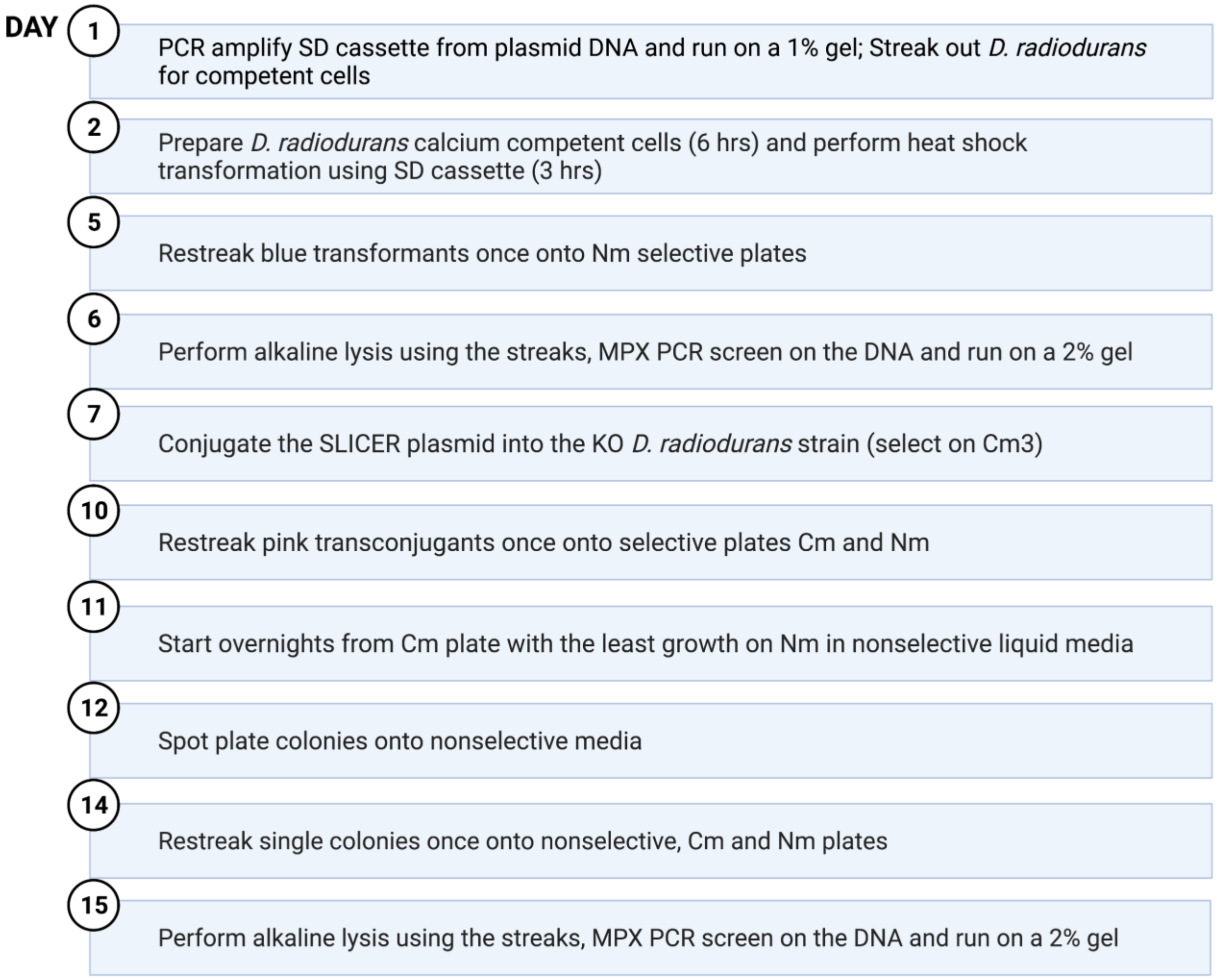


**Figure S2.** Step-by-step SLICER protocol. Laboratory protocol for the SLICER method, which can be used to create a seamless gene deletion in *D. radiodurans* in approximately 2 weeks. Created with BioRender.com.

### Supplemental Tables

**Table S1.** *Deinococcus radiodurans* strains created in this study.

| Strain | Description | Resistance | Reference or Source |
|---|---|---|---|
| ΔRM1 | ΔORF14075 | None | This study |
| ΔRM1-2 | ΔORF14075 Δ*Mrr* | None | This study |
| ΔRM1-3 | ΔORF14075 Δ*Mrr* ΔORF15360 | None | This study |
| ΔRM1-4 | ΔORF14075 Δ*Mrr* ΔORF15360 Δ*Mrr2* | None | This study |
| ΔRM1-5 Nm | ΔORF14075 Δ*Mrr* ΔORF15360 Δ*Mrr2* ΔORF2230 | Nm | This study |

**Table S2.** List of plasmids used in this study.

| Plasmid | Description | Resistance | Reference or Source |
|---|---|---|---|
| pBH474 | Suc$^s$ derivative of pTH474 | | [1] |
| pDEINO1 | Replicating plasmid with codon-optimized Cm marker | Cm (*D. radiodurans*, *E. coli*)<br>Nm (*D. radiodurans*, *E. coli*, *S. meliloti*)<br>HIS3 (*S. cerevisiae*)<br>Ntc (*P. tricornutum*) | [2]<br>Addgene ID: 179472 |
| pDEINO3 | Replicating plasmid with Tet marker | Tet and Cm (*D. radiodurans*, *E. coli*)<br>HIS3 (*S. cerevisiae*) | [2]<br>Addgene ID: 179487 |
| pDEINO4 | Replicating plasmid with Strep marker | Cm (*D. radiodurans*, *E. coli*)<br>Strep (*D. radiodurans*)<br>HIS3 (*S. cerevisiae*) | [2]<br>Addgene ID: 179488 |
| pDEINO10 | Nonreplicating plasmid with two 1 kb homology regions flanking ORF2230 | Nm (*D. radiodurans*), Cm (*E. coli*), HIS3 (*S. cerevisiae*) | [2] |
| pSLICER | Replicating SLICER plasmid containing *I-SceI* endonuclease | Cm (*D. radiodurans*, *E. coli*)<br>HIS3 (*S. cerevisiae*) | This study |
| pSD1 | Non-replicating plasmid containing RM1 SD cassette | Nm and Tet (*D. radiodurans*, *E. coli*)<br>Cm (*E. coli*)<br>HIS3 (*S. cerevisiae*) | This study |
| pSD2 | Non-replicating plasmid containing RM2 SD cassette | Nm (*D. radiodurans*, *E. coli*)<br>Strep (*D. radiodurans*)<br>Cm (*E. coli*)<br>HIS3 (*S. cerevisiae*) | This study |
| pSD3 | Non-replicating plasmid containing RM3 SD cassette | Nm and Tet (*D. radiodurans*, *E. coli*)<br>HIS3 (*S. cerevisiae*) | This study |
| pSD4 | Non-replicating plasmid containing RM4 SD cassette | Nm and Tet (*D. radiodurans*, *E. coli*) | This study |

| | | | |
|---|---|---|---|
| | | HIS3 (*S. cerevisiae*) | |
| pET-24α(+)-lacZ | pET-24α(+) with *lacZ* under a constitutive promoter | Kan (*E. coli*)<br>HIS3 (*S. cerevisiae*) | Pellegrino, unpublished |
| pRAD1 | General cloning vector for use in *E. coli* or *D. radiodurans* | Amp (*E. coli*), Cm (*D. radiodurans*) | [3] |
| pTA-Mob | Broad-host-range mobilization plasmid | Gm (*E. coli*) | [4] |

**Table S3.** List of oligonucleotides used in this study. The bold, underlined sequence in the assembly primers represents the binding portion of the primer, while the remainder of the sequence is the hook (*i.e.*, homology region) to the adjacent fragment. Sequence in red indicates the *I-Sce*I recognition site.

| Name | Sequence (5' to 3') | Description |
|---|---|---|
| | pSLICER Assembly Primers | |
| BK1388_F | AGTACATCACCGACGAGCAAGGCAAGACGATC TTAATTAA**TCGAGCTGGTTGCCCTCGCC** | pSLICER assembly primer (split pCC1BAC-yeast #1) |
| BK1388_R | GAAGAGCGTTGATCAATGGCCTGTTCAAAAAC AGTTCTCA**TCCGGATCTGACCTTTACCA** | pSLICER assembly primer (split pCC1BAC-yeast #1) |
| BK1389_F | GTACGTGAAACGGATGAAGTTGGTAAAGGTCA GATCCGGA**TGAGAACTGTTTTTGAACAG** | pSLICER assembly primer (split pCC1BAC-yeast #2) |
| BK1389_R | TCGATAGATCTCGAGGCCTCGCGAGCTTGGCG TAATCATG**GTTTAAACGGGCTTCGCCCT** | pSLICER assembly primer (split pCC1BAC-yeast #2) |
| BK1390_F | GCTCGCCGCAGTCGAGCGACAGGGCGAAGCCC GTTTAAAC**CATGATTACGCCAAGCTCGC** | pSLICER assembly primer (Drad origin) |
| BK1945_R | TGTCCAGGGCCCTCGGTCTCCATGGCCCTCAG GCCCTCGC**TTTAGCTTCCTTAGCTCCTG** | pSLICER assembly primer (Drad origin) |
| BK1946_F | CCGAGCTTCGACGAGATTTTCAGGAGCTAAGG AAGCTAAA**GCGAGGGCCTGAGGGCCATG** | pSLICER assembly primer (DrCm$^R$) |
| BK1946_R | GGCTTGATTTTCAGAATAGGGGCCAATCCAGA ATTACCTC**AAAAAACCCCCCGGATTGCC** | pSLICER assembly primer (DrCm$^R$) |
| BK1947_F | GCAGAAAAAATCCCCCCGGTGGCAATCCGGGG GGTTTTTT**GAGGTAATTCTGGATTGGCC** | pSLICER assembly primer (*I-Sce*I endonuclease) |
| BK1947_R | CGCTATAATGACCCCGAAGCAGGGTTATGCAG CGGAAGAT**GAGCAGAGGCTCTCGCTGAT** | pSLICER assembly primer (*I-Sce*I endonuclease) |
| BK1948_F | TGGCCCTCACCGCCGCGTCCATCAGCGAGAGC CTCTGCTC**ATCTTCCGCTGCATAACCCT** | pSLICER assembly primer (*oriT*) |
| BK1392_R | CGCCAGCCCAGCGGCGAGGGCAACCAGCTCGA TTAATTAA**GATCGTCTTGCCTTGCTCGT** | pSLICER assembly primer (*oriT*) |
| | pSD1 (RM1) Assembly Primers | |
| BK1388_F | AGTACATCACCGACGAGCAAGGCAAGACGATC TTAATTAA**TCGAGCTGGTTGCCCTCGCC** | pSD1 assembly primer (split pCC1BAC-yeast #1) |
| BK1388_R | GAAGAGCGTTGATCAATGGCCTGTTCAAAAAC AGTTCTCA**TCCGGATCTGACCTTTACCA** | pSD1 assembly primer (split pCC1BAC-yeast #1) |

| | | |
|---|---|---|
| BK1389_F | GTACGTGAAACGGATGAAGTTGGTAAAGGTCA<br>GATCCGGA**TGAGAACTGTTTTTGAACAG** | pSD1 assembly primer (split pCC1BAC-yeast #2) |
| BK2093_R | CACGGCCGCGCTCGGCCTCTCTGGCGGCCTTCT<br>GGCGCTC**GGGCTTCGCCCTGTCGCTCG** | pSD1 assembly primer (split pCC1BAC-yeast #2) |
| BK2094_F | CCAGTAGTGCTCGCCGCAGTCGAGCGACAGGG<br>CGAAGCCC**GAGCGCCAGAAGGCCGCCAG** | pSD1 assembly primer (Tet$^R$) |
| BK2094_R | GAGCGCAATGCCCCGATCACCTTGCGGTACCG<br>GGCGAGGC**GATCAGACGCTGAGTGCGCT** | pSD1 assembly primer (Tet$^R$) |
| BK2095_F | GCAGGACGCCGATGATTTGAAGCGCACTCAGC<br>GTCTGATC**GCCTCGCCCGGTACCGCAAG** | pSD1 assembly primer (ORF14075 homology #1) |
| BK2095_R | AGGGCAGTTGGAAAGTTGAGGAAAGCAGGCG<br>TGTGTACCAGGTGCCCCGCGACGTTGCGT**GTG**<br>**CGCAGGAGTGGGCCACA** | pSD1 assembly primer (ORF14075 homology #1) |
| BK2096_F | TCCTCAACTTTCCAACTGCCCTCTCCGACTGAC<br>TTTGCTGCT**TAGGGATAACAGGGTAAT****GGGG**<br>**TGGGCGAAGAACTCCA** | pSD1 assembly primer (Nm$^R$) |
| BK2096_R | TCTCATTTCACTAAATAATAGTGAACGGCAGG<br>TATATGTG**AGCTTCACGCTGCCGCAAGC** | pSD1 assembly primer (Nm$^R$) |
| BK2097_F | GCAGCCCTTGCGCCCTGAGTGCTTGCGGCAGC<br>GTGAAGCT**CACATATACCTGCCGTTCAC** | pSD1 assembly primer (*SacB*) |
| BK2097_R | AGGGCAGTTGGAAAGTTGAGGAAAGCAGGCG<br>TGTGTACCAGGTGCCCCGCGACGTTGCGT**GGC**<br>**CATCGGCATTTTCTTTT** | pSD1 assembly primer (*SacB*) |
| BK2098_F | TGGTACACACGCCTGCTTTCCTCAACTTTCCAA<br>CTGCCCTCTCCGACTGACTTTGCTGCT**AGCTTG**<br>**AGATTCGTACTCGC** | pSD1 assembly primer (ORF14075 homology #2) |
| BK2098_R | CGCTATAATGACCCCGAAGCAGGGTTATGCAG<br>CGGAAGAT**CCCAGACCTGCTCCGGCGTG** | pSD1 assembly primer (ORF14075 homology #2) |
| BK2099_F | CTGAAAAAAGCCCGCATCGGCACGCCGGAGCA<br>GGTCTGGG**ATCTTCCGCTGCATAACCCT** | pSD1 assembly primer (*oriT*) |
| BK1392_R | CGCCAGCCCAGCGGCGAGGGCAACCAGCTCGA<br>TTAATTAA**GATCGTCTTGCCTTGCTCGT** | pSD1 assembly primer (*oriT*) |
| | pSD2 (RM2) Assembly Primers | |
| BK1388_F | AGTACATCACCGACGAGCAAGGCAAGACGATC<br>TTAATTAA**TCGAGCTGGTTGCCCTCGCC** | pSD2 assembly primer (split pCC1BAC-yeast #1) |
| BK1388_R | GAAGAGCGTTGATCAATGGCCTGTTCAAAAAC<br>AGTTCTCA**TCCGGATCTGACCTTTACCA** | pSD2 assembly primer (split pCC1BAC-yeast #1) |
| BK1389_F | GTACGTGAAACGGATGAAGTTGGTAAAGGTCA<br>GATCCGGA**TGAGAACTGTTTTTGAACAG** | pSD2 assembly primer (split pCC1BAC-yeast #2) |
| BK2299_R | ACAAGCATAAAGCTTGCTCAATCAATCACCGG<br>ATCCCCGG**GGGCTTCGCCCTGTCGCTCG** | pSD2 assembly primer (split pCC1BAC-yeast #2) |
| BK2300_F | CCAGTAGTGCTCGCCGCAGTCGAGCGACAGGG<br>CGAAGCCC**CCGGGGATCCGGTGATTGAT** | pSD2 assembly primer (Spec$^R$) |
| BK2300_R | CCAGCGGCTACGGGCGATGTACGAGCAGGAAC<br>TGGGATTC**GGATCCGGTGATTGATTGAG** | pSD2 assembly primer (Spec$^R$) |
| BK2301_F | GTTTACAAGCATAAAGCTTGCTCAATCAATCA<br>CCGGATCC**GAATCCCAGTTCCTGCTCGT** | pSD2 assembly primer (*Mrr* Homology #1) |
| BK2301_R | GCTGGAGTTCTTCGCCCACCCC**ATTACCCTGT**<br>**TATCCCTA****ATTTTTTAAGTTTACGCTCT** | pSD2 assembly primer (*Mrr* Homology #1) |
| BK2302_F | ACAGAGCGTAAACTTAAAAAAT**TAGGGATAA**<br>**CAGGGTAAT****GGGGTGGGCGAAGAACTCCA** | pSD2 assembly primer (Nm$^R$) |

| | | |
|---|---|---|
| BK2302_R | TGTGAGCTAGCATTATACCTAGGACTGAGCTAGCTGTCAA**AGCTTCACGCTGCCGCAAGC** | pSD2 assembly primer (Nm$^{R}$) |
| BK2303_F | GCAGCCCTTGCGCCCTGAGTGCTTGCGGCAGCGTGAAGCT**TTGACAGCTAGCTCAGTCCT** | pSD2 assembly primer (*lacZ*) |
| BK2303_R | GTCTGGACTTACGGCTTTCGTCCCTTCCGCGCACCCAGCGCCTGTCCCAGCGACGCCCGC**TATAAACGCAGAAAGGCCCA** | pSD2 assembly primer (*lacZ*) |
| BK2304_F | CGCTGGGTGCGCGGAAGGGACGAAAGCCGTAAGTCCAGACAGAGCGTAAACTTAAAAAAT**GCTCGCCTGACAGGGCGGTT** | pSD2 assembly primer (*Mrr* Homology #2) |
| BK2304_R | CGCTATAATGACCCCGAAGCAGGGTTATGCAGCGGAAGAT**CGGCAGTTCCACGTTGCACA** | pSD2 assembly primer (*Mrr* Homology #2) |
| BK2305_F | CACCGCCCACATCGCCGAGTTGTGCAACGTGGAACTGCCG**ATCTTCCGCTGCATAACCCT** | pSD2 assembly primer (*oriT*) |
| BK1392_R | CGCCAGCCCAGCGGCGAGGGCAACCAGCTCGATTAATTAA**GATCGTCTTGCCTTGCTCGT** | pSD2 assembly primer (*oriT*) |
| | pSD3 (RM3) Assembly Primers | |
| BK2390_F | CAAAGGCCTGCACGTCCTCAAAGAGCAGCGGCTGAATCAC**ATCTTCCGCTGCATAACCCT** | pSD3 assembly primer (*oriT* + split pCC1BAC-yeast #1) |
| BK2092_R | GAAGAGCGTTGATCAATGGCCTGTTCAAAAACAGTTCTCA**TCCGGATCTGACCTTTACCA** | pSD3 assembly primer (*oriT* + split pCC1BAC-yeast #1) |
| BK2093_F | GTACGTGAAACGGATGAAGTTGGTAAAGGTCAGATCCGGA**TGAGAACTGTTTTTGAACAG** | pSD3 assembly primer (split pCC1BAC-yeast #2 +Tet$^{R}$) |
| BK2391_R | GGGCCTACATAAAAGGATCAGTCCTCGGAAACTTCTGCCC**GATCAGACGCTGAGTGCGCT** | pSD3 assembly primer (split pCC1BAC-yeast #2 +Tet$^{R}$) |
| BK2392_F | GCAGGACGCCGATGATTTGAAGCGCACTCAGCGTCTGATC**GGGCAGAAGTTTCCGAGGAC** | pSD3 assembly primer (ORF15360 Homology #1) |
| BK2392_R | GCTGGAGTTCTTCGCCCACCCCATTACCCTGTTATCCCTA**ACAATGCCATTTATGTTTTC** | pSD3 assembly primer (ORF15360 Homology #1) |
| BK2393_F | CAGAAAACATAAATGGCATTGTTAGGGATAACAGGGTAAT**GGGGTGGGCGAAGAACTCCA** | pSD3 assembly primer (Nm$^{R}$ + *lacZ*) |
| BK2395_R | TGGCACTTCGGCACTAGCTGCGTCAGCCTTGTTTATTGACTCCGGCACGACTTGGAGACG**TATAAACGCAGAAAGGCCCA** | pSD3 assembly primer (Nm$^{R}$ + *lacZ*) |
| BK2394_F | GTCAATAAACAAGGCTGACGCAGCTAGTGCCGAAGTGCCAGAAAACATAAATGGCATTGT**GCCCCCTCGACCTTGCCCGT** | pSD3 assembly primer (ORF15360 Homology #2) |
| BK2394_R | CGCTATAATGACCCCGAAGCAGGGTTATGCAGCGGAAGAT**GTGATTCAGCCGCTGCTCTT** | pSD3 assembly primer (ORF15360 Homology #2) |
| | pSD4 (RM4) Assembly Primers | |
| BK2408_F | GTTCGACCAAATGCGCCCCCCCACCCGCAGCGTCAGGTCG**ATCTTCCGCTGCATAACCCT** | pSD4 assembly primer (*oriT* + split pCC1BAC-yeast #1) |
| BK2092_R | GAAGAGCGTTGATCAATGGCCTGTTCAAAAACAGTTCTCA**TCCGGATCTGACCTTTACCA** | pSD4 assembly primer (*oriT* + split pCC1BAC-yeast #1) |
| BK2093_F | GTACGTGAAACGGATGAAGTTGGTAAAGGTCAGATCCGGA**TGAGAACTGTTTTTGAACAG** | pSD4 assembly primer (split pCC1BAC-yeast #2 +Tet$^{R}$) |
| BK2409_R | GCACACCCTGGTCGGCGCCGGGGCGACAGAGGGCGGCAGT**GATCAGACGCTGAGTGCGCT** | pSD4 assembly primer (split pCC1BAC-yeast #2 +Tet$^{R}$) |
| BK2410_F | GCAGGACGCCGATGATTTGAAGCGCACTCAGCGTCTGATC**ACTGCCGCCCTCTGTCGCCC** | pSD4 assembly primer (*Mrr2* Homology #1) |

| | | |
|---|---|---|
| BK2410_R | GCTGGAGTTCTTCGCCCACCCCATTACCCTGTTATCCCTA**<u>TCGCCCACCTGATGATCGAG</u>** | pSD4 assembly primer (*Mrr2* Homology #1) |
| BK2411_F | TACTCGATCATCAGGTGGGCGATAGGGATAACAGGGTAAT**<u>GGGGTGGGCGAAGAACTCCA</u>** | pSD4 assembly primer ($Nm^R$ + *lacZ*) |
| BK2413_R | TACGGCGTGGGCGTGCTGACCCGCGAGACCTACCAGATTCGCCGCTTAGACGCGGATTAT**<u>TATAAACGCAGAAAGGCCCA</u>** | pSD4 assembly primer ($Nm^R$ + *lacZ*) |
| BK2412_F | GAATCTGGTAGGTCTCGCGGGTCAGCACGCCCACGCCGTACTCGATCATCAGGTGGGCGA**<u>AATTCGGCCAGTCGCCGGTA</u>** | pSD4 assembly primer (*Mrr2* Homology #2) |
| BK2412_R | CGCTATAATGACCCCGAAGCAGGGTTATGCAGCGGAAGAT**<u>CGACCTGACGCTGCGGGTGG</u>** | pSD4 assembly primer (*Mrr2* Homology #2) |
| | Seamless Deletion Cassette Amplification | |
| BK1965_F | CCGGTACCGCAAGGTGAT | RM1 |
| BK1965_R | GGGTCGGTGTCCATCTCTT | RM1 |
| BK1964_F | CCCAGTTCCTGCTCGTACAT | RM2 |
| BK1964_R | AGTTCCACGTTGCACAACTC | RM2 |
| BK1966_F | GCAGAAGTTTCCGAGGACTG | RM3 |
| BK1966_R | GCTGCTCTTTGAGGACGTG | RM3 |
| BK2378_F | GGTGTGCAGCTCGTCTATGA | RM4 |
| BK2378_R | CGCATTTGGTCGAACAGC | RM4 |
| | Seamless Deletion Multiplex Primers | |
| BK2451_F | GAACGGGTGCAAATCAAGAC | Drad gDNA control – 150 bp |
| BK2451_R | CCGCGTCACCGAGTACAT | Drad gDNA control – 150 bp |
| BK2005_F | ACGACCATCACACCACTGAA | Plasmid backbone – 645 bp |
| BK2005_R | CATGACCAGCGTTTATGCAC | Plasmid backbone – 645 bp |
| BK2000_F | CGAAACGATCCTCATCCTGT | Nm marker – 311 bp |
| BK2000_R | AGGAAGCGGAACACGTAGAA | Nm marker – 311 bp |
| BK2003_F | GGCCCACTTCATCACAGAGT | RM1 (ORF14075) – 509 bp |
| BK2003_R | CCGAACAGGTCCTGGAAGTA | RM1 (ORF14075) – 509 bp |
| BK2216_F | CCTGACCGAAAGAGAGTTCG | RM2 (*Mrr*) – 508 bp |
| BK2216_R | GTAGCGGGAGGTCGTCATAA | RM2 (*Mrr*) – 508 bp |
| BK2004_F | GCTGGTAAATGCCCTTCGTA | RM3 (ORF15360) – 510 bp |
| BK2004_R | TCTACGCCGACTTCCTGTTC | RM3 (ORF15360) – 510 bp |
| BK2450_F | CTGAACCCGGACGTAGTGAT | RM4 (Mrr2) – 460 bp |
| BK2450_R | TTCAGACCATTCCGGCTTAC | RM4 (Mrr2) – 460 bp |

### Supplemental Methods

**Plasmid Design and Construction.** All plasmids in this study (Table S2) were constructed from PCR amplified DNA fragments assembled using a yeast spheroplast transformation method as previously described[5]. The primers used to amplify the fragments for plasmid assembly (Table S3) contained 20 bp binding and 40 bp of overlapping homology to the adjacent DNA fragment. Following assembly, DNA was isolated from *S. cerevisiae* and the plasmid pool was electroporated into *E. coli* Epi300. Plasmids from individual colonies were screened for correct assembly using multiplex PCR and diagnostic restriction digest. All plasmids were built to contain a pCC1BAC-yeast backbone allowing replication and selection in *E. coli* (chloramphenicol) and *S. cerevisiae* (-HIS) with a low-copy *E. coli* origin that can be induced to high copy with arabinose. They also have an origin of transfer (*oriT*) necessary for conjugation. pSD1-4: nonreplicating plasmids containing two ~1 kb regions of homology flanking ORF14075, *Mrr*, ORF15360, and *Mrr2*, respectively, amplified from wild-type *D. radiodurans* genomic DNA. Between the homology regions on the plasmids is an I-*Sce*I recognition site, a selective marker (*nptII*) and visual screening marker (*lacZ*) amplified from pDEINO1 and pET-24α(+)-lacZ, respectively, and an 80 bp duplication of the 3' end of homology region 1. The aforementioned elements make up the SD cassette. These plasmids also contain a second selective marker for *D. radiodurans* outside of the SD cassette, *tetR/A* or *aadA1* amplified from pDEINO3 and pDEINO4, respectively[2]. pSLICER: replicating plasmid built to contain a *D. radiodurans* codon-optimized *cat* gene under the control of a constitutive promoter (drKatA) and origin of replication amplified from pDEINO1[2]. A synthesized *D. radiodurans* codon-optimized I-*Sce*I endonuclease gene was also incorporated on this plasmid under the control of the PDR_2508 promoter and terminator set[6].

***D. radiodurans* Genomic DNA Isolation.** Alkaline lysis was performed using 3 mL of saturated culture as previously described[5] to extract *D. radiodurans* genomic DNA for analysis.

**Multiplex PCR Analysis of *D. radiodurans* Knockouts.** Multiplex PCR analysis was performed according to the manufacturer's instructions for "Standard Multiplex PCR" (Qiagen Multiplex PCR Handbook) with the following modifications and the primers listed in Table S3. A final volume of 20 μL was used and reaction mix components were adjusted accordingly. A volume of 1 μL of undiluted template DNA and 1 μL of dimethyl sulfoxide (DMSO) was used in the reaction mix. Thermocycler conditions were chosen according to the "Universal Multiplex Cycling Protocol" with the initial activation step decreased to 5 min, using an annealing temperature of 60°C, and 30 cycles. Gel electrophoresis was used to visualized 2 μL of the PCR product on a 2% agarose gel.

**Spot Plating *D. radiodurans***. *D. radiodurans* was grown overnight in 5 mL cultures of TGY media supplemented with the appropriate antibiotics (none, neomycin or chloramphenicol). The cultures were diluted to an $OD_{600}$ of 0.1 before performing 10-fold serial dilutions in TGY media up to $10^{-5}$ dilution. Then, 5 μL of each dilution was plated on nonselective TGY media and/or TGY media supplemented with appropriate antibiotics and incubated at 30°C for 2-3 days.

***D. radiodurans* Growth Curve and Doubling Time Calculation**. Growth rates were evaluated for *D. radiodurans* strains: wild type, ΔRM1, ΔRM1-2, ΔRM1-3, ΔRM1-4, and ΔRM1-5 $Nm^R$. Single colonies were inoculated into 5 mL of liquid TGY media and grown overnight at 30°C with shaking at 225 rpm. Cultures were diluted to an $OD_{600}$ of 0.1 in the same media, and 200 μL of each culture was aliquoted into a 96-well plate, along with a TGY media only control. In the Epoch 2 (BioTek, USA) plate reader, strains were grown at 30°C with continuous, orbital shaking (559 cpm). Absorbance ($A_{600}$) measurements were taken every 15 min for 24 h for a total of 97 readings using Gen5 data analysis software version 3.08.01 (Biotek, USA). This experiment was performed with three biological replicates, each with two technical replicates. Growth curves were plotted with data points representing the average of six measurements for each strain with error bars representing standard error of the mean. For simplicity, every other time point was omitted; therefore, readings are presented for every 30 min and the curve is cut off at the 17 hour time point when cultures approached end point density. The doubling time of each replicate was determined using the R package Growthcurver (Sprouffske K., Growthcurver, http://github.com/sprouffske/growthcurver, 2016)[7]. The doubling time is reported as an average of the six replicates for each strain, and the standard deviation was calculated.
